# Brassinosteroids Suppress Ethylene Biosynthesis via Transcription Factor BZR1 in Pear and Apple Fruit

**DOI:** 10.1101/2020.02.27.968800

**Authors:** Yinglin Ji, Yi Qu, Zhongyu Jiang, Xin Su, Pengtao Yue, Xinyue Li, Yanan Wang, Haidong Bu, Hui Yuan, Aide Wang

## Abstract

The plant hormone ethylene is important for the ripening of climacteric fruit, such as pear (*Pyrus ussuriensis*), and the brassinosteroid (BR) class of phytohormones affects ethylene biosynthesis during ripening, although via an unknown molecular mechanism. Here, we observed that exogenous BR treatment suppressed ethylene production during pear fruit ripening, and that the expression of the transcription factor *PuBZR1* was enhanced by epibrassinolide (EBR) treatment during pear fruit ripening. PuBZR1 was shown to interact with PuACO1, which converts 1-aminocyclopropane-1-carboxylic acid (ACC) to ethylene, and suppress its activity. We also observed that BR-activated PuBZR1 bound to the promoters of *PuACO1* and of *PuACS1a*, which encodes ACC synthase, and directly suppressed their transcription. Moreover, PuBZR1 suppressed the expression of transcription factor *PuERF2* by binding its promoter, and PuERF2 bound to the promoters of *PuACO1* and *PuACS1a*. We concluded that PuBZR1 indirectly suppresses the transcription of *PuACO1* and *PuACS1a* through its regulation of PuERF2. Ethylene production and the expression profiles of the corresponding apple (*Malus domestica*) homologs showed similar changes following EBR treatment. Together, these results suggest that BR-activated BZR1 suppresses ACO1 activity and the expression of *ACO1* and *ACS1a*, thereby reducing ethylene production during pear and apple fruit ripening. This likely represents a conserved mechanism by which exogenous BR suppresses ethylene biosynthesis during climacteric fruit ripening.

**One-sentence summary:** BR-activated BZR1 suppresses ACO1 activity and expression of *ACO1* and *ACS1a*, which encode two ethylene biosynthesis enzymes, thereby reducing ethylene production during pear and apple fruit ripening.

## INTRODUCTION

Fruit ripening is a plant developmental process that can be categorized as climacteric or non-climacteric (Klee and Giovannoni, 2011). The onset of normal ripening in climacteric fruit requires increased biosynthesis of the gaseous hormone ethylene (Barry and Giovannoni, 2007), and reducing ethylene in climacteric fruit leads to slow softening rate and longer shelf-life (Osorio et al., 2013). Numerous studies have shown that other hormones are also involved (Zaharah et al., 2011; Chai et al., 2012; Li et al., 2017), however, current understanding of the mechanisms by which these hormones interact with ethylene signaling to regulate fruit ripening is very limited.

The biosynthesis of ethylene includes two critical steps: the formation of 1-aminocyclopropane-1-carboxylic acid (ACC) from *S*-adenosyl methionine (SAM) by the enzyme ACC synthase (ACS) and the conversion of ACC to ethylene by ACC oxidase (ACO) (Yang and Hoffman, 1984). Previous reports have documented the importance of *ACS* and *ACO* genes in ethylene biosynthesis during fruit ripening. For example, silencing of *ACS* or *ACO* in transgenic tomato (*Solanum lycopersicum*) or apple (*Malus domestica*) fruit results in substantially reduced or undetectable ethylene production (Dandekar et al., 2004; Schaffer et al., 2007; Gupta et al., 2013). The actions of both *ACS* and *ACO* have been shown to be regulated transcriptionally in many species. Examples include a MADS-box transcription factor, *RIPENING INHIBITOR* (*RIN*), which binds to the promoter of *SlACS2* in tomato (Ito et al., 2008), an ethylene response factor, MaERF11, which binds to the promoter of *MaACO1* and suppresses its expression in banana (*Musa acuminata*) (Han et al., 2016), and MdERF3, which binds to the promoter of *MdACS1* and activates its expression in apple (Li et al., 2016).

Various phytohormones have been observed to influence ethylene biosynthesis during fruit ripening. Abscisic acid (ABA) concentration increases at the onset of tomato fruit ripening and application of exogenous ABA promotes the expression of ethylene biosynthetic genes and ethylene production (Zhang et al., 2009). In peach (*Prunus persica*) fruit, the level of the auxin, indole-3-acetic acid (IAA) increases prior to fruit ripening and the application of synthetic auxin has been reported to result in increased *PpACS1* expression and ethylene production (Trainotti et al., 2007; Tatsuki et al., 2013). Another well-studied example is jasmonate, which promotes ethylene production in apple fruit via the MdMYC2 transcription factor. This process includes jasmonate-activated MdMYC2 binding to the promoters of both *MdACS1* and *MdACO1* to induce their expression during fruit ripening (Li et al., 2017). However, in contrast to the above classes of phytohormones, the mechanism by which brassinosteroids (BRs) affect ethylene biosynthesis during ripening is not known.

BRs are involved in regulating a wide range of plant physiological processes and much has been learnt about the BR signaling pathway (Clouse, 2011). Following biosynthesis, BRs are perceived by BRASSINOSTEROID INSENSITIVE 1 (BRI1), leading to association with BRI1-associated kinase 1 (BAK1). BRI1 and BAK1 transphosphorylate each other, allowing BRI1 to phosphorylate BR SIGNALING KINASE 1 (BSK1). The phosphorylated BSK1 activates BRI SUPPRESSOR 1 (BSU1), which inhibits BRASSINOSTEROID INSENSITIVE 2 (BIN2) by dephosphorylation, leading to accumulation of unphosphorylated BRASSINAZOLE-RESISTANT 1 (BZR1) and its homologs in the nucleus. BZR1 and its homologs bind to the promoters of BR-responsive genes and regulate their expression (He et al., 2005; Yin et al., 2005; Li and Jin, 2007; Kim and Wang, 2010; Clouse, 2011). The effect of BR on ethylene biosynthesis and fruit ripening has been documented. For example, BR-treated jujube (*Zizyphus jujuba*) fruit, which is categorized as a climacteric fruit, shows significantly reduced ethylene production during storage (Zhu et al., 2010), while strawberry (*Fragaria ananassa*), a non-climacteric fruit, shows delayed fruit ripening after application of epibrassinolide (EBR) (Chai et al., 2012). In tomato, another climacteric fruit, treatment with brassinolide promotes the expression of *SlACS* and *SlACO* genes, as well as ethylene production (Zhu et al., 2015). More interestingly, overexpression of a BR biosynthetic gene *DWARF* in tomato results in increased level of endogenous BR and ethylene production, and earlier ripening (Li et al., 2016), indicating endogenous BR can affect fruit ripening.

These studies indicate that BR is involved in the regulation of ethylene biosynthesis and fruit ripening; however, little is known about how BR signaling genes interact with ethylene biosynthetic genes to regulate ethylene production. In this study, we investigated the effects of exogenous BR on fruit ripening in pear (*Pyrus ussuriensis*) and apple. The resulting data provide new insights into the molecular basis by which BR suppresses ethylene biosynthesis during climacteric fruit ripening.

## RESULTS

### BR Inhibits Ethylene Production in Pear Fruit

To investigate the effect of BR on ethylene biosynthesis during climacteric fruit ripening, we sampled pear (*P. ussuriensis*) fruit at commercial harvest in 2015 and 2016. Fruit sampled in each year were treated with EBR, a brassinosteroid, and stored at room temperature for 15 days (d) (Fig. 1A). Following EBR treatment, ethylene production was significantly lower and fruit firmness was significantly higher compared with untreated control fruit (Fig. 1B; Supplemental Fig. S1A-C).

**Fig. 1.**
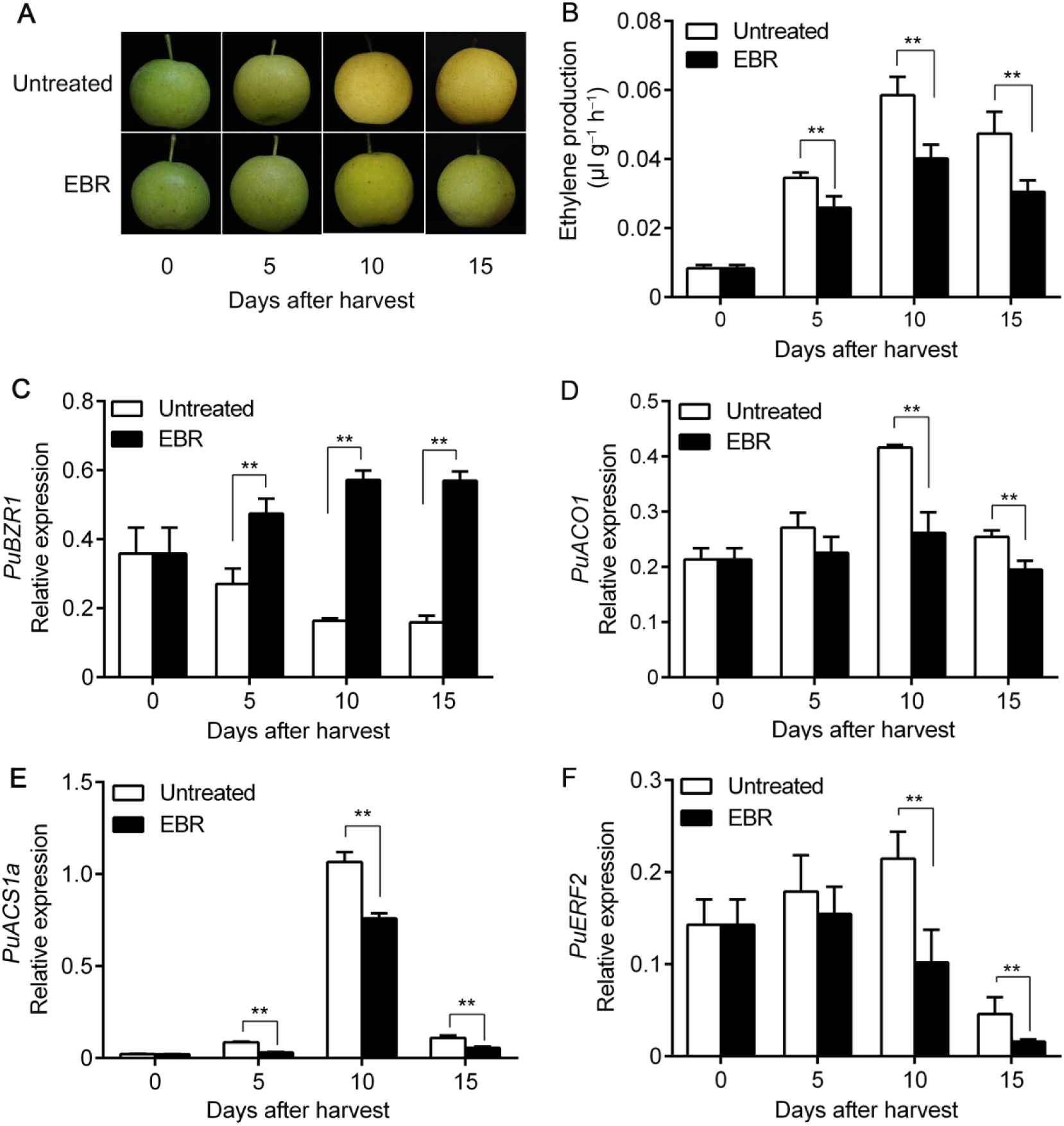
Ethylene production and gene expression in pear fruit treated with epibrassinolide (EBR). Pear fruit were collected at commercial harvest in 2016, treated with EBR, and stored at room temperature for 15 days **(A)**. Ethylene production was measured **(B)**, and the expression levels of *PuBZR1* **(C)**, *PuACO1* **(D)**, *PuACS1a* **(E)** and *PuERF2* **(F)** were investigated by quantitative reverse transcription (qRT)-PCR. Untreated, fruit not receiving any treatment; EBR, fruit treated with EBR. The *x*-axes indicate the number of days of storage at room temperature after harvest. Three biological replicates were analyzed as described in the **Methods** section. Values represent means ± SE. Statistical significance was determined using a Student’s *t-*test (\*\**P*<0.01).

Given that BZR1 is a key transcription factor in the BR signaling pathway (Kim and Wang, 2010), as an initial step in understanding BR regulated processes associated with ethylene production and fruit ripening, we identified a total of seven *PuBZR1* or *PuBZR1-like* genes from pear (Supplemental Fig. S2). Of these, we observed that only the expression of *PuBZR1* was significantly enhanced by EBR treatment (Fig. 1C; Supplemental Fig. S2). We therefore focused on *PuBZR1* and tested the hypothesis that it acts as a BR-induced suppressor of ethylene biosynthesis during pear fruit ripening.

### PuBZR1 Interacts with PuACO1 and Suppresses PuACO1 Enzyme Activity

To further characterize the putative role of *PuBZR1* in BR-suppressed ethylene biosynthesis, we used PuBZR1 as a bait in a yeast two-hybrid (Y2H) screen of a pear fruit cDNA library. A total of 135 positive clones were identified from the screen, corresponding to 23 genes; one of which encoded PuACO1. The potential interaction between PuBZR1 and PuACO1 was then confirmed by co-expressing the two proteins in yeast cells (Supplemental Fig. S3A), and this was further validated using a pull-down assay involving PuBZR1-His and PuACO1-GST peptide tagged fusion proteins (Supplemental Fig. S3B). We divided the predicted coding region of *PuBZR1* into two and the cording region of *PuACO1* into four fragments, and used them in a Y2H assay, which showed that the PuBZR1 N terminal region (PuBZR1N) interacts with both the N- and D fragments of PuACO1 (PuACO1N and PuACO1D) (Fig. 2A). Interestingly, the PuACO1D contains Fe^2+^ binding sites (Supplemental Fig. S4) that is essential for ACO enzyme activity (Shaw et al., 1996; Zhang et al., 1997; Rocklin et al., 1999). We then investigated the intracellular localization of PuBZR1 and PuACO1. The coding sequence (CDS) of PuBZR1 and PuACO1 fused to a green fluorescent protein (GFP) peptide tag were infiltrated into tobacco (*Nicotiana benthamiana*) leaves. The result showed that PuBZR1 and PuACO1 localized in cytoplasm and nucleus (Supplemental Fig. 3C). Next, we investigated whether the interaction between PuBZR1 and PuACO1 affected PuACO1 enzymatic activity, in reactions containing purified PuACO1 mixed with different amounts of purified PuBZR1. PuACO1 activity gradually declined with increasing amounts of PuBZR1 (Fig. 2B), suggesting suppression of PuACO1 activity through direct interaction with PuBZR1. We also measured ACO enzyme activity in extracts from pear fruit that had been treated with EBR, or were untreated, and found that EBR treatment significantly inhibited ACO activity (Supplemental Fig. S5).

**Fig. 2.**
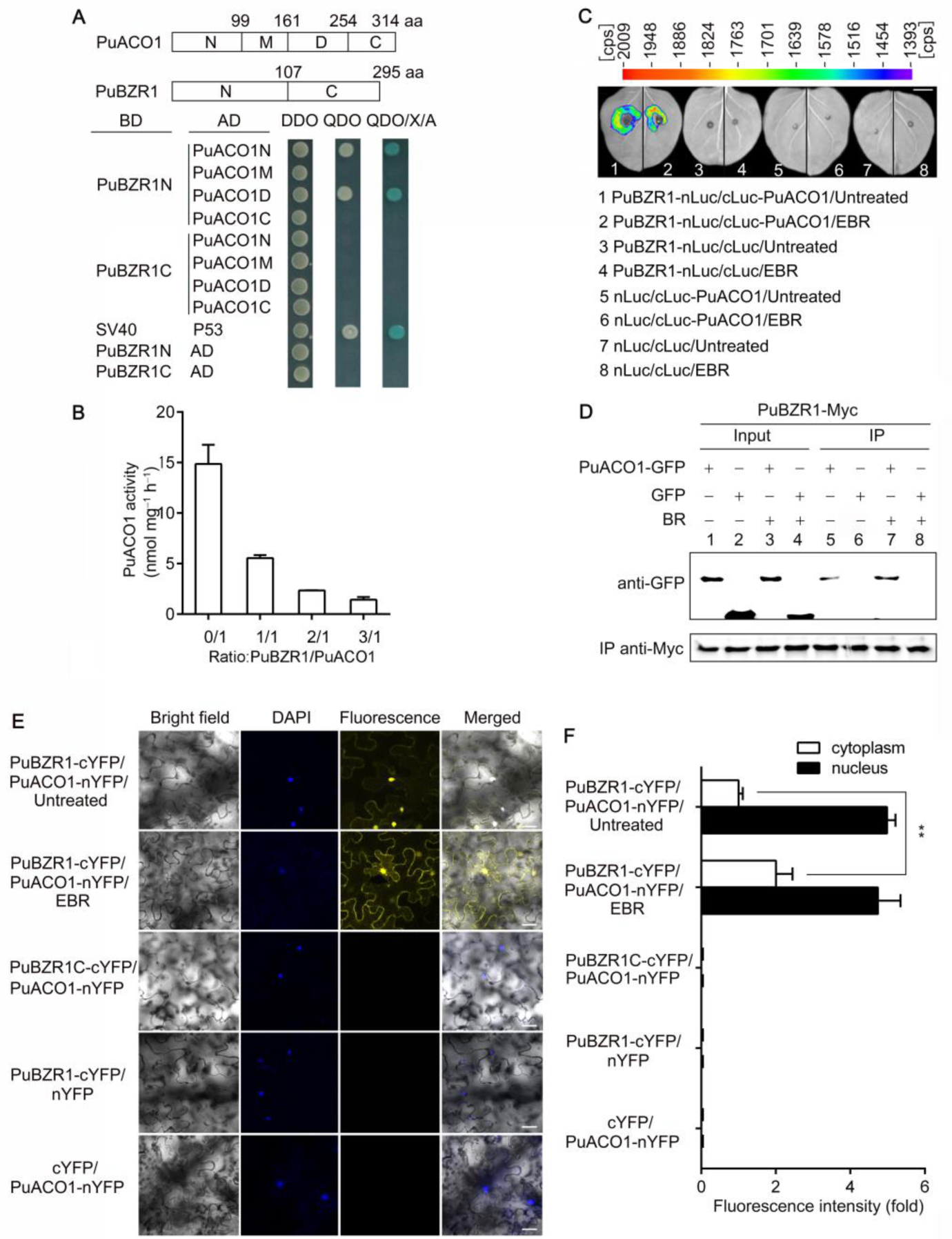
Brassinosteroid (BR)-activated PuBZR1 interacts with PuACO1, which inhibits PuACO1 enzyme activity. **(A)** The PuBZR1 and PuACO1 protein sequences were divided into two and four fragments, respectively, and their interaction was investigated using a yeast two-hybrid assay. DDO, SD medium lacking Trp and Leu; QDO, SD medium lacking Trp, Leu, His and Ade; QDO/X/A, QDO medium containing x-a-gal and aureobasidin A. SV40 and P53 were used as a positive control, and AD vectors as negative controls. The blue color indicates protein interaction. **(B)** The influence of PuBZR1-PuACO1 interaction on PuACO1 activity was evaluated by adding increased amounts of the PuBZR1 protein. Recombinant His-tagged PuBZR1 and GST-tagged PuACO1 were used. Error bars represent the SE of three independent measurements. **(C)** A firefly luciferase complementation imaging assay showing that epibrassinolide (EBR) treatment enhanced the interaction between PuBZR1 and PuACO1 in tobacco leaves. *Agrobacterium tumefaciens* strain EHA105 harboring different constructs was infiltrated into tobacco leaves. Untreated, tobacco leaves not receiving any treatment; EBR, tobacco leaves treated with EBR. Luciferase activities were recorded in these regions 3 d after infiltration. Bar, 1 cm; cps, signal counts per second. **(D)** A coimmunoprecipitation (co-IP) assay showing that epibrassinolide (EBR) treatment enhanced the interaction between PuBZR1 and PuACO1 in tobacco leaves. PuBZR1 fused to a Myc tag (PuBZR1-Myc) and PuACO1 fused to a GFP tag (PuACO1-GFP) was overexpressed in tobacco leaves and a Myc antibody was used for immunoprecipitation analysis. Myc and GFP antibodies were used in an immunoblot analysis. The band detected by the GFP antibody in the precipitated protein sample indicates the interaction between PuBZR1 and PuACO1 (lane 5) and EBR treatment enhances the interaction (lane 7). **(E)** Interaction of PuBZR1 and PuACO1 in a bimolecular fluorescence complementation assay. Tobacco leaves were co-infiltrated with PuBZR1-cYFP and PuACO1-nYFP constructs and visualized by confocal microscopy 48 h after infiltration. EBR treatment was applied to the infiltrated tobacco leaves 3 h before imaging. DAPI (2-(4-Amidinophenyl)-6-indolecarbamidine dihydrochloride) was used as a nuclear marker. PuBZR1C-cYFP with PuACO1-nYFP, PuBZR1-cYFP with nYFP, and cYFP with PuACO1-nYFP, were used as negative controls. Scale bars, 20 μM. **(F)** Fluorescence intensity of BiFC in cytoplasm in (E). The fluorescence signal was quantified using Image J software from 10 randomly selected cytoplasm or nucleus regions of each treatment. Values represent means ± SE. Statistical significance was determined using a Student’s *t-*test (\*\**P*<0.01).

**Fig. 3.**
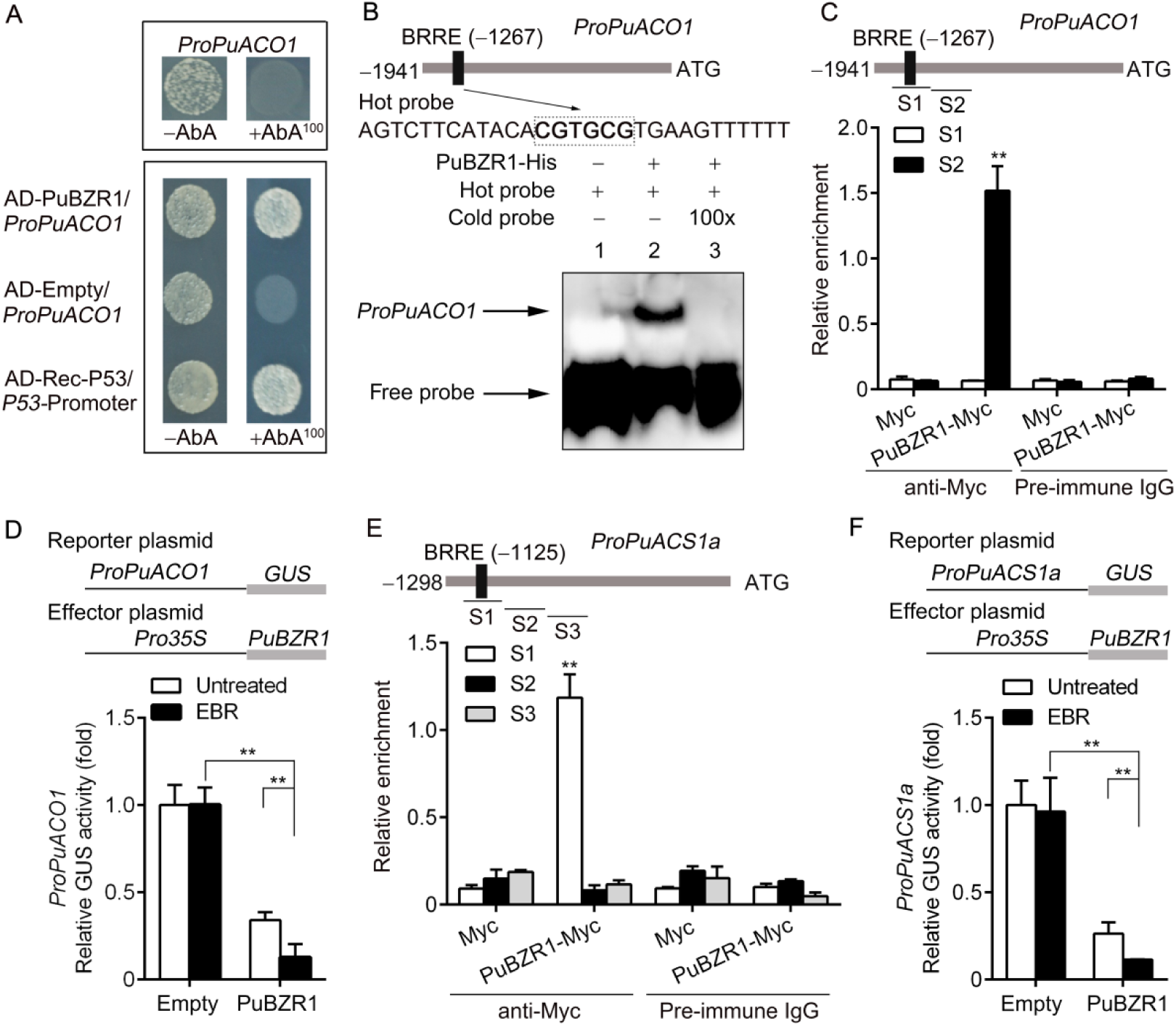
BR-Activated PuBZR1 suppresses the transcription of *PuACO1* and *PuACS1a*. **(A)** Yeast one-hybrid (Y1H) analysis showing that PuBZR1 binds to the promoter of *PuACO1* (*ProPuACO1*). AbA (aureobasidin A), a yeast cell growth inhibitor, was used as a screening marker. The basal concentration of AbA was 100 ng ml^-1^. Rec-P53 and the *P53*-promoter were used as positive controls. The empty vector and the *PuACO1* promoter were used as negative controls. **(B)** Electrophoretic mobility shift assay (EMSA) showing that PuBZR1 binds to the BRRE motif in the *PuACO1* promoter. The hot probe was a biotin-labeled fragment of the *PuACO1* promoter containing the BRRE motif, and the cold probe was a non-labeled competitive probe (100-fold that of the hot probe). His-tagged PuBZR1 was purified. **(C)** Chromatin Immunoprecipitation (ChIP)-PCR showing the *in vivo* binding of PuBZR1 to the *PuACO1* promoter. Cross-linked chromatin samples were extracted from PuBZR1-Myc overexpressing pear fruit calli (PuBZR1-Myc) and precipitated with an anti-Myc antibody. Eluted DNA was used to amplify the sequences neighboring the BRRE motif by quantitative (q)-PCR. Two regions (S1 and S2) were investigated. Fruit calli overexpressing the Myc sequence (Myc) were used as negative controls. Values are the percentage of DNA fragments that were co-immunoprecipitated with the Myc antibody or a non-specific antibody (pre-immune rabbit IgG) relative to the input DNA. The ChIP assay was repeated three times and the enriched DNA fragments in each ChIP were used as one biological replicate for qPCR. Values represent means ± SE. Asterisks indicate significantly different values (\*\**P*<0.01). **(D)** β-Glucuronidase (GUS) activity analysis showing that PuBZR1 suppresses the *PuACO1* promoter. The PuBZR1 effector vector and the reporter vector containing the *PuACO1* promoter were infiltrated into tobacco leaves to analyze the regulation of GUS activity. Untreated, tobacco leaves not receiving any treatment; EBR, tobacco leaves treated with epibrassinolide. Three independent transfection experiments were performed. Values represent means ± SE. Asterisks indicate significantly different values (\*\**P*<0.01). **(E)** ChIP-PCR showing the *in vivo* binding of PuBZR1 to the *PuACS1a* promoter. **(F)** GUS activity analysis showing that PuBZR1 suppresses the *PuACS1a* promoter.

**Fig. 4.**
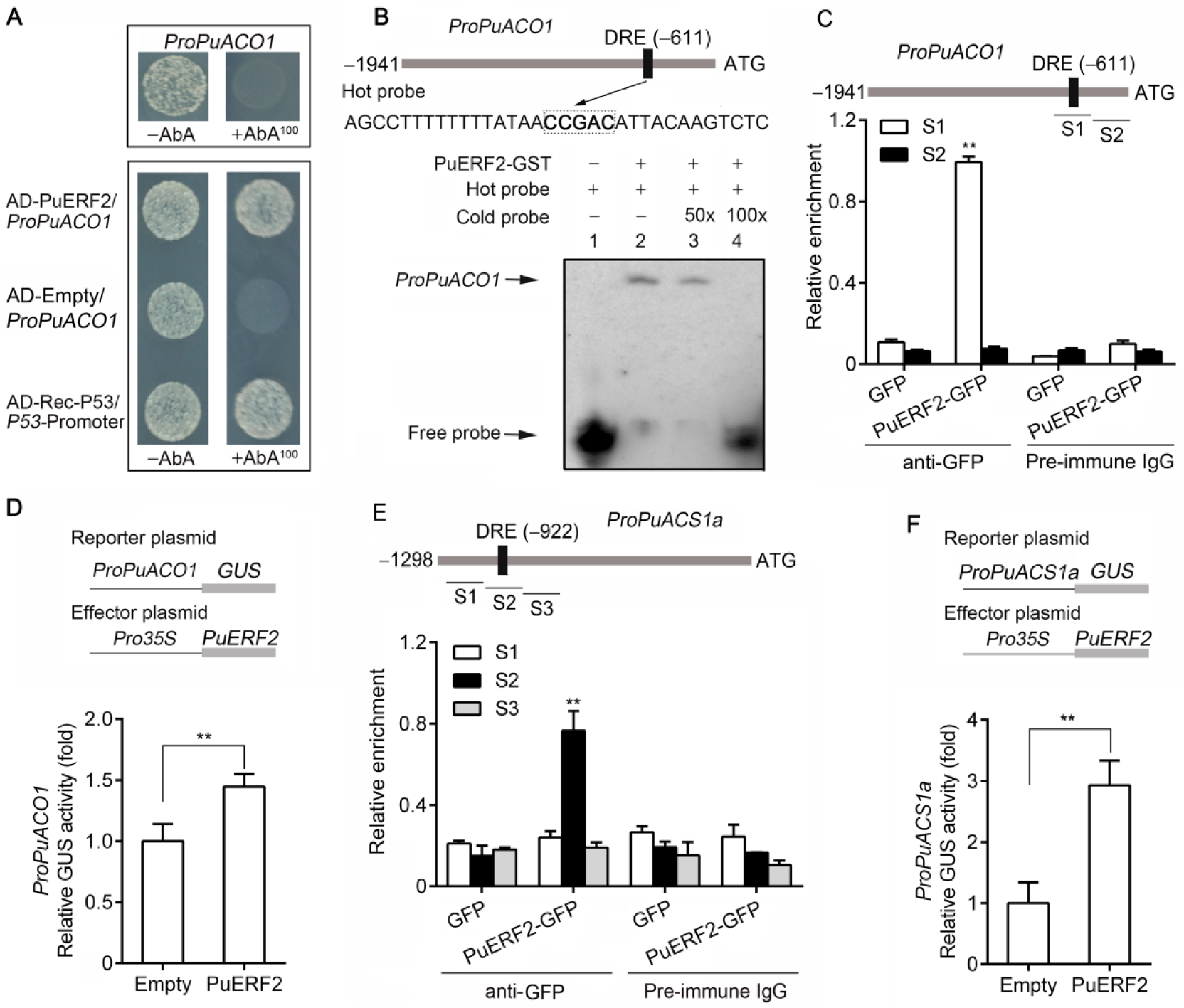
PuERF2 suppresses *PuACO1* transcription. **(A)** Yeast one-hybrid (Y1H) analysis showing that PuERF2 binds to the promoter of *PuACO1* (*ProPuACO1*). **(B)** Electrophoretic mobility shift assay (EMSA) showing that PuERF2 binds to the DRE motif of the *PuACO1* promoter. The hot probe was a biotin-labeled fragment of the *PuACO1* promoter containing the DRE motif, and the cold probe was a non-labeled competitive probe (50- and 100-fold that of the hot probe). GST-tagged PuERF2 was purified. **(C)** Chromatin Immunoprecipitation (ChIP)-PCR showing the *in vivo* binding of PuERF2 to the *PuACO1* promoter. Cross-linked chromatin samples were extracted from PuERF2-green fluorescent protein (GFP) overexpressing fruit calli (PuERF2-GFP) and precipitated with an anti-GFP antibody. Eluted DNA was used to amplify the sequences neighboring the DRE motif by quantitative (q)-PCR. Two regions (S1 and S2) were investigated. Fruit calli overexpressing the GFP sequence (GFP) were used as negative controls. Values are the percentage of DNA fragments that co-immunoprecipitated with the GFP antibody or a non-specific antibody (pre-immune rabbit IgG) relative to the input DNA. The ChIP assay was repeated three times and the enriched DNA fragments in each ChIP were used as one biological replicate for qPCR. Values represent means ± SE. Asterisks indicate significantly different values (\*\**P*<0.01). **(D)** β-Glucuronidase (GUS) activity analysis showing that PuERF2 promotes the activity of the *PuACO1* promoter. The PuERF2 effector vector and the reporter vector containing the *PuACO1* promoter were infiltrated into tobacco leaves to analyze the regulation of GUS activity. Three independent transfection experiments were performed. Values represent means ± SE. Asterisks indicate significantly different values (\*\**P*<0.01). **(E)** ChIP-PCR showing the *in vivo* binding of PuERF2 to the *PuACS1a* promoter. **(F)** GUS activity analysis showing that PuERF2 promotes the *PuACS1a* promoter.

Next, we tested whether EBR treatment affects the PuBZR1/PuACO1 interaction using a firefly luciferase (Luc) complementation imaging assay. Constructs containing PuBZR1 fused with the N terminus of Luc (PuBZR1-nLuc) and the C terminus of Luc fused with PuACO1 (cLuc-PuACO1) were co-infiltrated into tobacco leaves, and the infiltrated leaves were treated with EBR. A luminescence signal was detected in the PuBZR1-nLuc/cLuc-PuACO1 co-expressing region (Fig. 2C, region 1) but not in the negative controls (Fig. 2C, regions 3, 5 and 7), consistent with a PuBZR1/PuACO1 protein interaction *in planta*. Following EBR treatment, a stronger luminescence signal was observed in the PuBZR1-nLuc/cLuc-PuACO1 co-expressing region (Fig. 2C, region 2) but not in the negative controls (Fig. 2C, regions 4, 6 and 8), indicating that EBR treatment enhances the interaction between PuBZR1 and PuACO1. A coimmunoprecipitation (co-IP) assay confirmed this result (Fig. 2D). Finally, a bimolecular fluorescence complementation (BiFC) assay was performed. Constructs containing PuBZR1 fused into pSPYCE-35S vector (PuBZR1-cYFP) and PuACO1 fused into pSPYNE-35S vector (PuACO1-nYFP) were co-infiltrated into tobacco leaves, and the infiltrated leaves were treated with EBR. The result showed that PuBZR1 interacted with PuACO1 in both cytoplasm and nucleus (Fig. 2E). Following EBR treatment, a stronger YFP signal was observed in cytoplasm (Fig. 2E and 2F), while no significant difference of the YFP signal was observed in nucleus (Fig. 2E and 2F). These results suggested that EBR treatment enhances the interaction between PuBZR1 and PuACO1 in cytoplasm.

### BR-Activated PuBZR1 Suppresses the Expression of *PuACO1* and *PuACS1a* via Transcriptional Regulation

We observed that *PuACO1* expression was reduced by EBR treatment (Fig. 1D), consistent with transcriptional regulation. We identified a BR response element (BRRE) in the promoter of *PuACO1* (1,941 bp upstream of the predicted translation start site) that we predicted might be involved in BZR1 binding, and so investigated whether PuBZR1 can bind to the *PuACO1* promoter and regulate its expression. This was indeed confirmed using both a yeast one-hybrid (Y1H) assay (Fig. 3A) and an electrophoretic mobility shift assay (EMSA) (Fig. 3B).

*In vivo* verification was performed by conducting a chromatin immunoprecipitation (ChIP)-PCR assay. The CDS of *PuBZR1* fused to a sequence encoding a Myc peptide tag was overexpressed in pear fruit calli. The presence of PuBZR1 substantially enhanced the PCR-based detection of the *PuACO1* promoter (Fig. 3C), indicating that PuBZR1 binds to the *PuACO1* promoter *in vivo*. When the regulation of the *PuACO1* promoter by PuBZR1 was examined in tobacco leaves co-transformed with the *Pro35S:PuBZR1* and *ProPuACO1:GUS* constructs using a β-glucuronidase (GUS) activation assay, a significantly reduced GUS signal was observed (Fig. 3D), indicating that PuBZR1 suppresses the activity of the *PuACO1* promoter. When EBR was applied to the tobacco leaves, the GUS signal was further reduced (Fig. 3D). Taken together, these results suggested that PuBZR1 directly suppresses the transcription of *PuACO1* and that BR strengthens this suppression.

ACS is also central to ethylene biosynthesis through its role in forming the ethylene precursor, ACC (Yang and Hoffman, 1984; Kende, 1993), and our previous study revealed five *ACS* genes that were differentially expressed during pear fruit ripening (Huang et al., 2014). When we investigated their expression profiles in EBR treated fruit in this current study, we detected high expression of *PuACS1a*, which was significantly suppressed by the EBR treatment (Fig. 1E; Supplemental Fig. S6). Notably, a BRRE motif was identified in the *PuACS1a* promoter (1,298 bp). We then performed ChIP-PCR and a GUS activation assay in tobacco, which revealed that PuBZR1 bound and reduced the promoter activity of *PuACS1a* (Fig. 3E and 3F). When EBR was applied to the tobacco leaves, the GUS signal was reduced further (Fig. 3F), suggesting that PuBZR1 directly inhibits the transcription of *PuACS1a*, and that BR strengthens this suppression.

### BR-Activated PuBZR1 Suppresses the Expression of *PuERF2*, and PuERF2 Binds to the *PuACO1* and *PuACS1a* Promoters

Our previous study showed that four ERF (ethylene response factor) transcription factors were differentially expressed during pear fruit ripening (Huang et al., 2014). Here, we investigated their expression and found that, of the four, only *PuERF2* expression was significantly suppressed by the EBR treatment (Fig. 1F), while the others showed no significant change compared with the controls (Supplemental Fig. 7). We next confirmed that PuERF2 can bind to the *PuACO1* promoter using both Y1H and EMSA analysis (Fig. 4A and 4B). We further demonstrated binding *in vivo* by ChIP-PCR, where the CDS of PuERF2 fused to a sequence encoding GFP peptide tag was overexpressed in pear fruit calli. The presence of PuERF2 substantially enhanced the PCR-based detection of the *PuACO1* promoter (Fig. 4C), indicating that PuERF2 binds to the *PuACO1* promoter *in vivo*. The regulation by PuERF2 of the *PuACO1* promoter was examined in tobacco leaves and we observed that PuERF2 activated the *PuACO1* promoter (Fig. 4D). We tested whether *PuACS1a* expression was transcriptionally regulated by PuERF2 and found, by ChIP-PCR and a GUS activation assay, that PuERF2 bound to and activated the *PuACS1a* promoter (Fig. 4E and 4F).

Given the presence of a BRRE motif in *PuERF2* promoter (1,979 bp), we investigated whether PuBZR1 can bind the *PuERF2* promoter and regulate its expression. Binding was indeed confirmed using both a Y1H assay (Fig. 5A) and an EMSA analysis (Fig. 5B), while a ChIP-PCR assay demonstrated that PuBZR1 can bind to the *PuERF2* promoter *in vivo* (Fig. 5C). The regulation of the *PuERF2* promoter by PuBZR1 was examined in tobacco leaves, and we determined that PuBZR1 suppresses the activity of the *PuERF2* promoter, while BR strengthens this suppression (Fig. 5D). These results suggest that BR-activated PuBZR1 indirectly suppresses the expression of *PuACO1* and *PuACS1a* through transcriptional regulation of PuERF2.

**Fig. 5.**
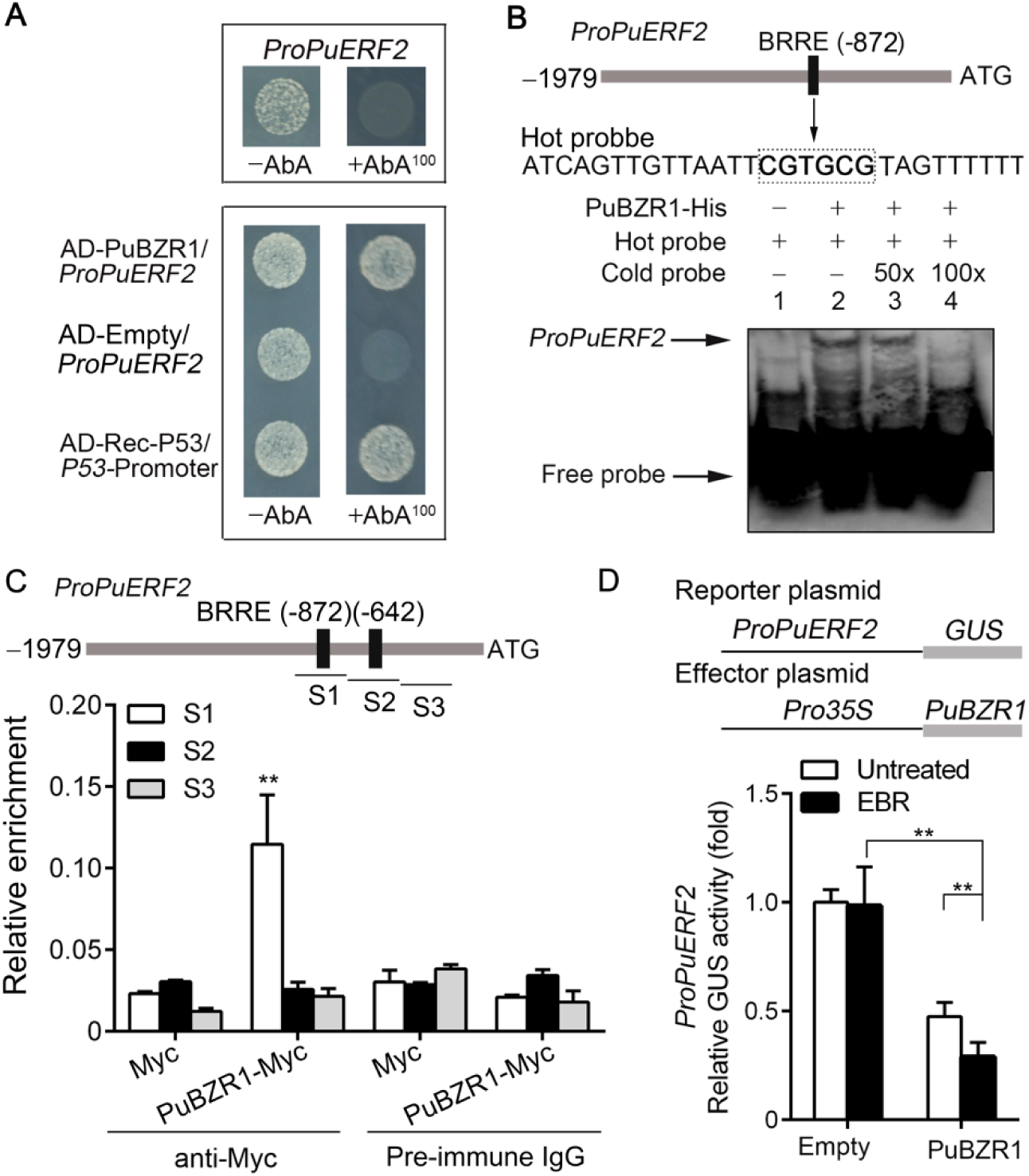
Brassinosteroid (BR)-activated PuBZR1 suppresses *PuERF2* transcription. **(A)** Yeast one-hybrid (Y1H) analysis showing that PuBZR1 binds to the promoter of *PuERF2* (*ProPuERF2*). **(B)** Electrophoretic mobility shift assay (EMSA) showing that PuBZR1 binds to the BRRE motif of the *PuERF2* promoter. The hot probe was a biotin-labeled fragment of the *PuERF2* promoter containing the BRRE motif, and the cold probe was a non-labeled competitive probe (50- and 100-fold that of the hot probe). His-tagged PuBZR1 was purified. **(C)** Chromatin Immunoprecipitation (ChIP)-PCR showing the *in vivo* binding of PuBZR1 to the *PuERF2* promoter. Cross-linked chromatin samples were extracted from PuBZR1-Myc overexpressing pear fruit calli (PuBZR1-Myc) and precipitated with an anti-Myc antibody. Eluted DNA was used to amplify the sequences neighboring the BRRE motif by qPCR. Three regions (S1-S3) were investigated. Fruit calli overexpressing the Myc sequence (Myc) were used as negative controls. Values are the percentage of DNA fragments that co-immunoprecipitated with the Myc antibody or a non-specific antibody (pre-immune rabbit IgG) relative to the input DNA. The ChIP assay was repeated three times and the enriched DNA fragments in each ChIP were used as one biological replicate for qPCR. Values represent means ± SE. Asterisks indicate significantly different values (\*\**P*<0.01). **(D)** β-Glucuronidase (GUS) activity analysis showing that PuBZR1 suppresses the *PuERF2* promoter. The PuBZR1 effector vector and the reporter vector containing the *PuERF2* promoter were infiltrated into tobacco leaves to analyze the regulation of GUS activity. Untreated, tobacco leaves not receiving any treatment; EBR, tobacco leaves treated with epibrassinolide. Three independent transfection experiments were performed. Values represent means ± SE. Asterisks indicate significantly different values (\*\**P*<0.01).

Next, we investigated whether PuERF2 can bind to the promoter of *PuBZR1 in vivo*, since it contains a DRE motif, but we found evidence through ChIP-PCR that it does not (Supplemental Fig. S8A). We also showed in tobacco leaves that PuERF2 regulation of the *PuACO1* and *PuACS1a* promoters was not EBR dependent (Supplemental Fig. S8B and 8C), suggesting that PuERF2 does not respond to EBR treatment without the presence of PuBZR1. We concluded that PuERF2 likely does not regulate the transcription of *PuBZR1*, and that PuBZR1 works upstream of PuERF2 in response to BR. We also investigated the potential interaction between PuBZR1 and PuERF2 using a Y2H assay, and determined that they do not interact with each other (Supplemental Fig. S9). Given the interaction between PuBZR1 and PuACO1 in the nucleus (Fig. 2E), we investigated the influence of this interaction on the binding of PuBZR1 to its target promoters. EMSA analyses were performed and the results showed that PuACO1 did not influence the binding of PuBZR1 to the promoters of *PuACO1, PuACS1a* and *PuERF2* (Supplemental Fig. S10).

### PuBZR1 Plays a Significant Role in BR-Suppressed Ethylene Production

To further confirm the role of *PuBZR1* in BR-suppressed ethylene biosynthesis, we transiently silenced *PuBZR1* expression in pear fruit. The full *PuBZR1* CDS was ligated into the pRI101 vector in the reverse direction and the resulting construct introduced into *Agrobacterium tumefaciens*, cultures of which were infiltrated into fruit still attached to trees. Fruit infiltrated with the empty pRI101 vector were used as a control. The infiltrated fruit were harvested at 7 DAI (days after infiltration), treated with EBR and stored at room temperature for 10 d (Fig. 6A). In the *PuBZR1*-suppressed pear fruit (PuBZR1-AN), PuBZR1 transcript levels were significantly reduced (Fig. 6B), and after EBR treatment they showed significantly higher ethylene production compared with the control fruit (Fig. 6C). In addition, the expression levels of *PuACO1, PuACS1a* and *PuERF2* (Fig. 6D-F), and enzyme activity of ACC oxidase (Fig. 6G) were higher than in control fruit. These findings indicate that PuBZR1 action is important for BR-suppressed ethylene production in pear fruit.

**Fig. 6.**
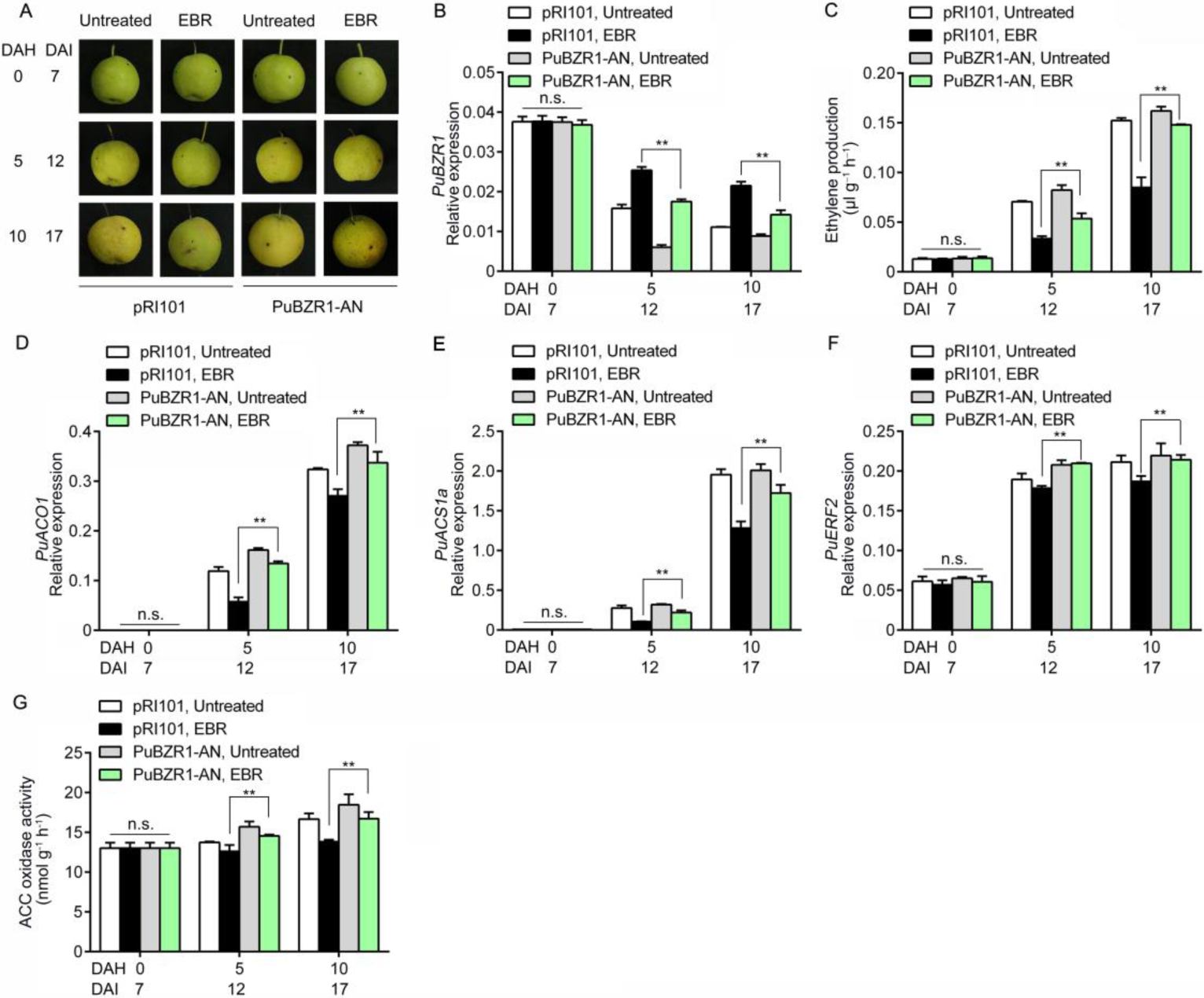
*PuBZR1* is involved in brassinosteroid (BR)-suppressed ethylene biosynthesis in pear fruit. *PuBZR1* was silenced in pear fruit (PuBZR1-AN) by *Agrobacterium tumefaciens-*mediated transient transformation. Fruit infiltrated with an empty pRI101 vector were used as controls (pRI101). PuBZR1-AN and control fruits were harvested 7 days after infiltration, treated with epibrassinolide (EBR) immediately and then stored at room temperature for 10 d **(A)**. *PuBZR1* expression was examined by quantitative reverse transcription (qRT)-PCR **(B)**. Ethylene production (**C**), the expression levels of *PuACO1* (**D**), *PuACS1a* (**E**) and *PuERF2* (**F**), and the ACO enzyme activity (**G**) were investigated. Untreated, fruit not receiving any treatment; EBR, fruit treated with EBR; DAH, days after harvest; DAI, days after infiltration. For qRT-PCR, three biological replicates were analyzed as described in the **Methods** section. Values represent means ± SE. Statistical significance was determined using a Student’s *t-*test (\*\**P*<0.01). n.s., no significant difference.

### BR also Suppresses Ethylene Production and the Expression of Ethylene Biosynthetic genes in Apple Fruit

To complement the studies in pear, we investigated the influence of BR on ethylene biosynthesis during ripening of apple (*M. domestica*) fruit, which are also climacteric. We sampled apple fruit at the commercial harvest stage and treated them with EBR. Fruit were then stored at room temperature for 25 d (Fig. 7A). Ethylene production in apple fruit treated with EBR was significantly lower and fruit firmness was significantly higher compared with those of the control fruit (Fig. 7B; Supplemental Fig. S1D). We compared the transcriptome of apple fruit stored at room temperature for 10 d and treated with or without EBR using RNA sequencing (RNA-seq). The transcript level of *MdBZR1* was increased, and that of the ethylene biosynthetic genes *MdACO1* and *MdACS1* was decreased as a consequence of EBR treatment (Supplemental Fig. S11; Supplemental Data Set 1). The qRT-PCR detected expression of these genes confirmed the result of RNA-seq (Fig. 7C-E). Moreover, we observed from the RNA-seq data that the expression of four *ERF* transcription factors was suppressed by the EBR treatment (Supplemental Data Set 2). Taken together, these results suggest that the mechanism for BR-suppressed ethylene biosynthesis in apple and pear fruit may be conserved.

## DISCUSSION

BR has been reported to participate in various aspects of plant development (Kim and Wang, 2010); however, although many studies have described the involvement of BR in fruit ripening (Clouse, 2011; Li et al., 2016; Baghel et al., 2019), the mechanism by which BR influences ethylene biosynthesis during this process is still not well understood. Here, we investigated the regulatory role of *PuBZR1* in BR-regulated ethylene biosynthesis during pear fruit ripening.

BR has been reported to be involved in fruit ripening in various species, including tomato (Zhu et al., 2015; Li et al., 2016), persimmon (*Diospyros kaki*) (He et al., 2018), mango (*Mangifera indica*) (Zaharah et al., 2011), jujube (Zhu et al., 2010), strawberry (Chai et al., 2012) and grape (*Vitis vinifera*) berry (Symons et al., 2006). However, these studies only investigated changes in ethylene production after BR treatment and the expression profile of genes involved in ethylene biosynthesis and signal transduction. In our study, we dissected the regulatory network involving PuBZR1 association with ethylene signaling genes in BR-suppressed ethylene production. We observed that BR-activated PuBZR1 binds to the *PuACO1* and *PuACS1a* promoters, directly down-regulating their expression (Fig. 3). A recent study demonstrated that MaBZR1 bound the promoters of *MaACS1* and *MaACO13/14* and repressed their expression in banana fruit (Guo et al., 2019), and our results support this model. In addition, we showed that PuBZR1 interacts with PuACO1 and suppresses its enzyme activity (Fig. 2), and that PuBZR1 binds to the *PuERF2* promoter and also down-regulates its expression (Fig. 5), while PuERF2 in turn binds to the promoters of *PuACO1* and *PuACS1a* (Fig. 4). Thus, PuBZR1 indirectly suppresses the expression of *PuACO1* and *PuACS1a* through PuERF2. These findings indicate a new direction for study the function of a transcription factor.

Our ChIP-PCR analysis showed no evidence of binding of PuERF2 to the *PuBZR1* promoter, despite the presence a predicted ERF binding site in the *PuBZR1* promoter (Supplemental Fig. S8A). A similar case was reported in a previous study in which MdERF3 showed no binding to its own promoter, although it contains a DRE motif in its promoter (Li et al., 2016). In addition, the PuERF2 regulation of the *PuACO1* and *PuACS1a* promoters was not affected by the EBR treatment in a GUS activation assay (Supplemental Fig. S8B and 8C), but the promoter of *PuERF2* responded to EBR treatment under the regulation of PuBZR1 (Fig. 5D). Taken together with the result that silencing *PuBZR1* expression in pear fruit suppressed the effect of the EBR treatment on ethylene production (Fig. 6), we conclude that PuBZR1 works upstream of PuERF2 plays a key role in BR-suppressed ethylene biosynthesis.

We observed that PuBZR1 suppressed PuACO1 activity by directly interacting with the PuACO1 protein (Fig. 2). ACO is a member of the Fe^2+^ dependent family of oxidases or oxygenases (Zhang et al., 1997) and it requires Fe^2+^ as a cofactor to catalyze the formation of ethylene (Dong et al., 1992). The ACO amino acid sequences are highly conserved between many species, and H177, D179 and H234 in ACOs of tomato, apple and avocado (*Persea americanna*) have been shown to be Fe^2+^ binding sites that are essential for enzyme activity (Shaw et al., 1996; Zhang et al., 1997; Rocklin et al., 1999). In these studies, substitutions of H177, D179 and H234 by site-directed mutagenesis resulted in complete loss of ACO activity. In our study, PuBZR1 interacted with the D fragment of PuACO1, which contains the three Fe^2+^ binding sites (Fig. 2; Supplemental Fig. S4). Moreover, EBR treatment enhanced the interaction between PuBZR1 and PuACO1 in the cytoplasm (Fig. 2), where ACO converts ACC to ethylene (Guy and Kende, 1984). Therefore, we propose that BR-enhanced PuBZR1/PuACO1 interaction might hinder the binding of Fe^2+^ to PuACO1, thereby suppressing PuACO1 activity.

Some studies have revealed that exogenous BR can promote ethylene production and accelerate fruit ripening in tomato (Vardhini and Rao, 2002; Zhu et al., 2015), persimmon (He et al., 2018) and mango (Zaharah et al., 2011). Although these fruits are also climacteric, the results of our studies of pear and apple fruits were opposite to these previous reports. This might be due to differences in species or the dose of BR applied: Zhu et al. (2010) reported that 5 μM of brassinolide suppressed ethylene production and fruit ripening in jujube, also a climacteric fruit, while application of 10 μM of brassinolide had the opposite result. In banana, application of 1, 2 or 4 μM of brassinolide accelerated the fruit ripening (Guo et al., 2019). In tomato fruit, 3 μM EBR promoted ethylene production and ripening (Vardhini and Rao, 2002), while in our study, 3 μM EBR inhibited ethylene production and ripening in both pear and apple fruit (Fig. 1 and Fig. 7), and a 10 μM EBR treatment had the same effect (data not shown). In strawberry and grape berry, both categorized as non-climacteric fruit, application of BR accelerated fruit ripening (Symons et al., 2006; Chai et al., 2012). In *Arabidopsis thaliana* seedlings, low concentration (10 or 100 nM) of exogenous BR can suppress ethylene biosynthesis, while high concentration (greater than 500 nM) of it promotes ethylene biosynthesis (Lv et al., 2018). These findings suggest that the influence of BR on ethylene biosynthesis and fruit ripening is different between species and might vary in a dose dependent manner.

Although we did not dissect the details of MdBZR1 regulation of MdACO1 or MdACS1 activity in apple, we observed that the EBR treatment also resulted in reduced ethylene production, reduced expression of *MdACO1* and *MdACS1*, and increased expression of *MdBZR1* in apple fruit (Fig. 7). Moreover, four ERF transcription factors down-regulated by an EBR treatment were identified from RNA-seq data (Supplemental Data Set 2). Thus the mechanism by which BR suppresses ethylene biosynthesis in apple fruit is likely similar to that in pear fruit.

In conclusion, BR-activated BZR1 suppressed ethylene biosynthesis during fruit ripening via three routes: (i) BZR1 suppressed the enzyme activity of ACO1 by direct protein interactions; (ii) BZR1 directly suppressed the transcription of *ACO1* and *ACS1a* by promoter binding; (iii) BZR1 indirectly suppressed the transcription of *ACO1* and *ACS1a* through *ERF2*, which bound the *ACO1* and *ACS1a* promoters (Fig. 8).

**Fig. 7.**
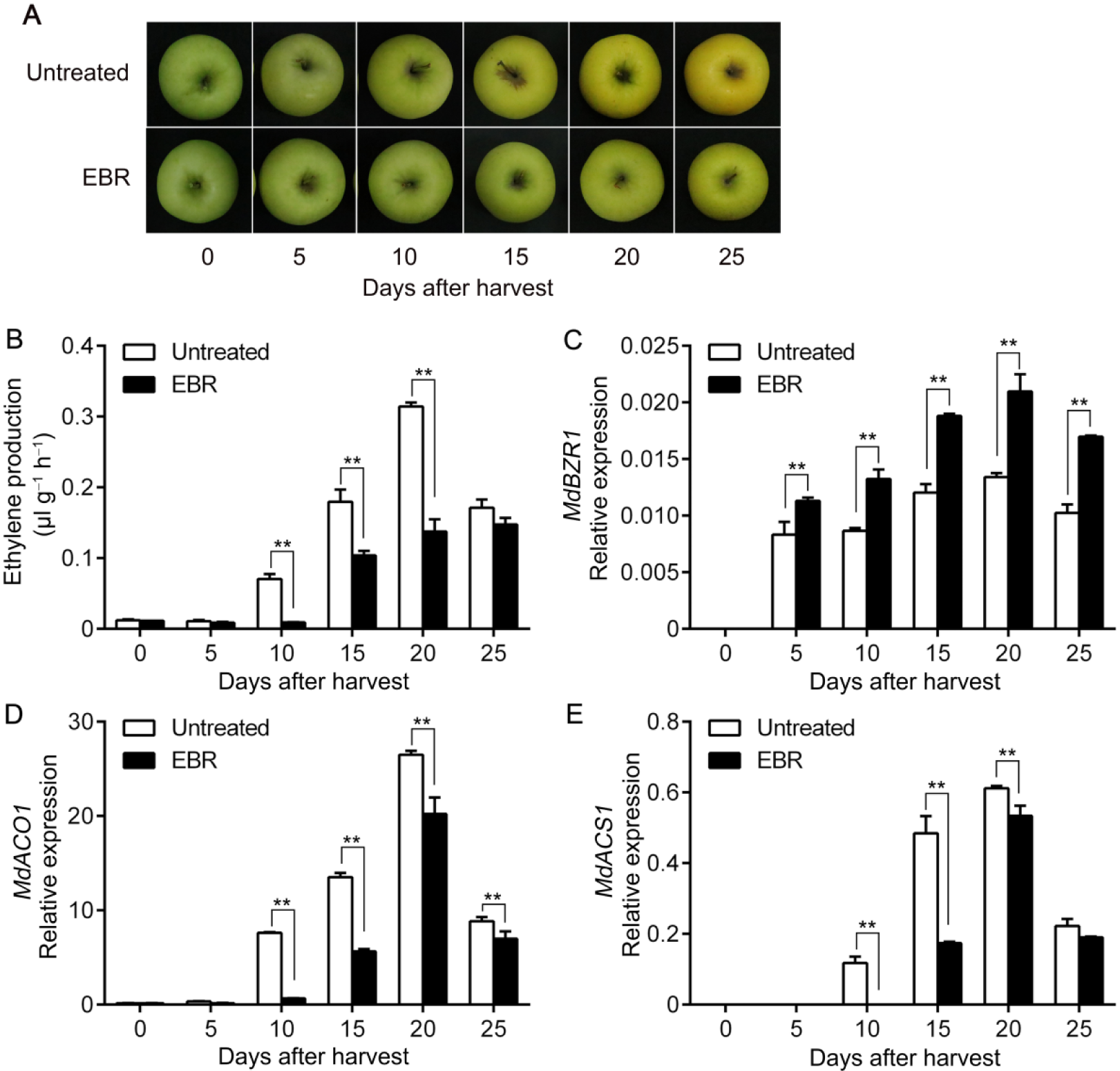
Ethylene production and gene expression in apple fruit treated with epibrassinolide (EBR). Apple fruit were collected at commercial harvest in 2017, treated with epibrassinolide (EBR), and then stored at room temperature for 25 d **(A)**. Ethylene production was measured **(B)**, and the expression levels of *MdBZR1* **(C)**, *MdACO1* **(D)** and *MdACS1* **(E)** were investigated by quantitative reverse transcription (qRT)-PCR. Untreated, fruit not receiving any treatment; EBR, fruit treated with EBR. Numbers under the *x*-axis indicate the number of days of storage at room temperature after harvest. Three biological replicates were analyzed as described in the **Methods** section. Values represent means ± SE. Statistical significance was determined using a Student’s *t-*test (\*\**P*<0.01).

**Fig. 8.**
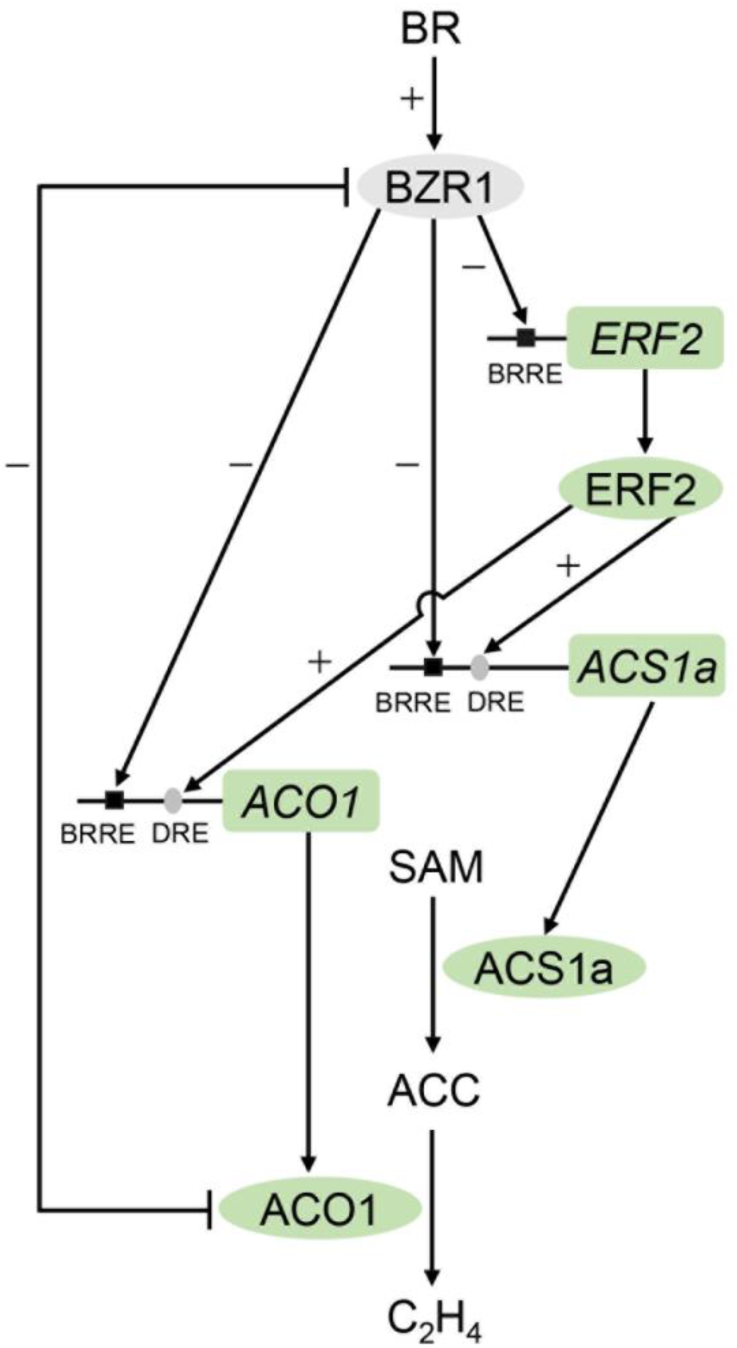
A model showing the suppression of ethylene biosynthesis by brassinosteroids (BR) through BZR1. BR-activated BZR1 interacts with ACO1, which inhibits ACO1 enzyme activity. The transcription factor, BZR1, binds to the promoter of *ACO1* and directly suppresses its transcription. BZR1 suppresses the activity of ERF2, which binds to the promoters of *ACO1* and *ACS1a*, and enhances their transcription: thus BZR1 suppresses *ACO1* and *ACS1a* transcription indirectly through ERF2. Through these three mechanisms BZR1 suppresses ACO1 enzyme activity and the transcription of *ACO1* and *ACS1a* to reduce ethylene biosynthesis in response to BR during pear fruit ripening. “+” promotion; “−”, suppression; BRRE, BR response element, BZR1-binding site; DRE, dehydration-responsive element, ERF-binding site; SAM, *S*-adenosyl methionine; ACC, 1-aminocyclopropane-1-carboxylic acid; C_2_H_4_, ethylene; BR, brassinosteroid.

## MATERIALS AND METHODS

### Plant Materials and Treatments

Pear (*P. ussuriensis* cv. Nanguo) fruit were obtained from the experimental farm of Shenyang Agricultural University. Fruit were collected on the day of commercial harvest, September 9 in 2015 and September 2 in 2016, when the content of total soluble solids reached to 12%. The fruit were divided into two groups (36 fruit per group). For the BR treatment, fruit in the first group were immersed in 3 μM epibrassinolide (EBR, Cat. no. 78821-43-9, Yuanye Biotechnology, Shanghai, China) for 2 hours. Fruit in the second group were not received any treatment and used as a control. After the treatments, the fruit were stored at room temperature for 15 d and sampled every 5 d. At each sampling time, 9 fruit were randomly collected and divided into 3 groups (3 fruit per group) resulting in three biological replicates. Ethylene production was measured as previously described (Li et al., 2014), and a total of three individual replicates were assayed. Statistical significance was determined using a Student’s *t*-test. Following ethylene measurement, flesh of the 3 fruit from each group was sliced, pooled, frozen in liquid nitrogen, and stored at each group was used as one biological replicate for gene expression analysis, and total of three replicates were used.

Apple (*M. domestica* cv. Golden Delicious) fruit were obtained from the experimental farm of Liaoning Pomology Institute (Xiongyue, China). Fruit were collected on the day of commercial harvest (September 26, 2017), treated with 3 μM EBR as above. Fruit not received any treatment were used as a control. Control and EBR treated fruits were stored at room temperature for 25 d and sampled every 5 d. The sampling regime was similar to that of the pear fruit. At each sampling time, 9 fruit were divided into 3 groups (as three biological replicates) for ethylene production measurement. Statistical significance was determined using a Student’s *t*-test. Following ethylene measurement, flesh from each group of fruit was pooled for RNA extraction and gene expression analysis, and a total of three replicates were used.

Pear (*P. ussuriensis* cv. Nanguo) fruit calli were prepared as previously described (Alayón-Luaces et al., 2008). Briefly, pear fruit harvested at day 75 after full bloom were used to generate primary calli, which were cultivated on basal Murashige and Skoog (MS) medium supplemented with 2 mg l^-1^ 2,4-dichlorophenoxyacetic acid (2,4-D, Sangon Biotech, http://www.life-biotech.com) and 1.5 mg l^-1^ 6-benzyladenine (6-BA, Sangon Biotech). Calli were subcultured on a proliferation medium consisting of basal MS medium supplemented with 2.5 mg l^-1^ 2,4-D and 1 mg l^-1^ 6-BA.

### Gene Expression Analysis

Total RNA was extracted according to the method of Li et al. (2014), and cDNA synthesis and quantitative reverse transcription PCR (qRT-PCR) were performed as previously described (Li et al., 2017). qRT-PCR was conducted using an Analytik Jena qTOWER^3^ G PCR System. RNA extracted from each group of flesh (as described above) was used as one biological replicate, and a total of three biological replicates were conducted. Statistical significance was determined using a Student’s *t*-test. Specific primers (Supplemental Data Set 3) for each gene were designed using Primer3 (http://frodo.wi.mit.edu). The pear and apple *Actin* genes (*PuActin* and *MdActin*, respectively) were used as internal controls.

### Yeast Two-Hybrid Assay

A cDNA library was constructed with mRNA from pear (*P. ussuriensis* cv. Nanguo) fruit harvested at commercial maturity in 2015, using a Make Your Own Mate & Plate Library System (Cat. no. 630489, Clontech). The *PuBZR1* CDS was introduced into the pGBKT7 vector enclosed in this kit using *Eco*RI and *Bam*HI sites. The recombinant plasmid was used as bait to screen the cDNA library using the Matchmaker™ Gold Yeast Two-Hybrid Library Screening System kit (Cat. no. 630489, Clontech).

The *PuACO1* (314 amino acids, aa), *PuACO1N* (aa 1-99), *PuACO1M* (aa 100-161), *PuACO1D* (aa 162-254) and *PuACO1C* (aa 255-314) sequences were introduced into the activation domain (AD) vector (pGADT7) using the *Nde*I and *Eco*RI restriction sites. The *PuBZR1* (295 aa), *PuBZR1N* (aa 1-107) and *PuBZR1C* (aa 108-295) sequences were ligated to the binding domain (BD) in the pGBKT7 vector using the *Nde*I and *Eco*RI restriction sites. The BD and AD vectors were co-transformed into the Y2HGold yeast strain. The detection of protein interactions between two proteins was conducted using the Matchmaker™ Gold Yeast Two-Hybrid Library Screening System kit.

### Protein Expression and Purification

The *PuBZR1* CDS was inserted into the pEASY-E1 vector (Transgen Biotech, http://www.transgen.com.cn) resulting in its downstream fusion to a His tag sequence. The *PuACO1* or *PuERF2* CDSs were inserted into the pGEX4T-1 vector (GE Healthcare, http://www3.gehealthcare.com) downstream from GST. The resulting plasmids were transformed into *Escherichia coli* BL21 (DE3) competent cells. Recombinant fusion proteins were purified as described in Li et al. (2016).

### Pull-Down Assay

To confirm the interaction between PuBZR1 and PuACO1, 5 μg of purified His fusion protein (PuBZR1-His) was bound to Ni-NTA His binding resin (Novagen). GST fusion proteins containing PuACO1 (PuACO1-GST) were added and incubated for 1 h at 4 °C with the subsequent immunoblot analysis performed as previously described (Li et al., 2017). GST protein was used as the negative control.

### Co-IP Assay

For the co-IP assay, the *PuBZR1* CDS was ligated into the pCAMBIA1307 vector (BioVector, http://www.biovector.net) to allow expression of the PuBZR1 protein with a Myc tag driven by the *CaMV 35S* promoter, using the *Xba*I and *Bam*HI sites. The CDS of PuACO1 was cloned into the *Kpn*I and *Eco*RI sites downstream of the GFP sequence and the *CaMV 35S* promoter in the pRI101 vector (TaKaRa). The recombinant *Pro35S:Myc-PuBZR1* and *Pro35S:GFP-PuACO1* constructs were infiltrated into tobacco (*N. benthamiana*) leaves using *Agrobacterium*-infiltration as previously described (Li et al., 2017), and EBR treatment (10 μM) was applied to the infiltrated leaves 3 h before sampling. Protein was extracted from the infiltrated tobacco leaves with or without EBR treatment and used for co-IP analysis. A Pierce coimmunoprecipitation kit (catalog no. 26149; Thermo Scientific) was used to immunoprecipitate Myc-PuBZR1 using 10 μl of anti-Myc antibody (1 mg ml^-1^; Transgen Biotech). The precipitate was analyzed by immunoblot analysis with the anti-GFP antibody (1 mg ml^-1^; Transgen Biotech) diluted 1:3000.

### Subcellular Localization

The *PuACO1* or *PuBZR1* coding region was cloned into the *Kpn*I and *Eco*RI sites downstream of GFP in the pRI101 vector to form the *Pro35S*:*GFP-PuACO1* or *Pro35S*:*GFP-PuBZR1* construct. The construct was co-infiltrated with a mCherry-labeled nuclear marker NF-YA4-mcherry (Zhang et al., 2019) into tobacco (*N. benthamiana*) leaves using *Agrobacterium*-infiltration. The tobacco plants were kept in the dark for 48 h after infiltration and then GFP fluorescence was observed under a confocal microscope (TCS SP8, Leica). *Pro35S:GFP* was used as a control. All transient expression assays were repeated at least three times, and the representative results were shown.

### BiFC Assay

The *PuACO1* CDS was ligated into the pSPYNE-35S vector (BioVector) using the *Xba*I and *Sal*I sites. The *PuBZR1* CDS was ligated into the pSPYCE-35S vector using the *Bam*HI and *Xho*I sites. The resulting plasmids were introduced into *Agrobacterium tumefaciens* strain EHA105, and then infiltration of wild tobacco leaves was performed. Infected leaves were analyzed 48 h after infiltration. EBR treatment (10 μM) was applied to the infiltrated tobacco leaves 3 h before imaging. YFP and 2-(4-Amidinophenyl)-6-indolecarbamidine dihydrochloride (DAPI, Beyotime Biotechnology, https://www.beyotime.com) fluorescence were observed under a confocal laser scanning microscope (TCS SP8, Leica). Fragments of PuBZR1C and PuACO1 were used as a negative control. All transient expression assays were repeated at least three times, and the representative results were shown. The quantitation of fluorescence signal was calculated from 10 randomly selected regions of each treatment using Image J software.

### Measurements of ACO Activity

PuACO1 enzyme activity was measured as previously described (Zhang et al., 1997). Purified PuACO1-GST protein (0.2 μg) was added to 2 ml of incubation buffer (pH 7.2) containing 10% (v/v) glycerol (Solarbio, http://www.solarbio.com), 5 mM Na-ascorbate (Sangon Biotech), 0.1 mM ACC (Sigma-Aldrich), 80 μM FeSO_4_ (Sangon Biotech), 15 mM NaHCO_3_ (Sangon Biotech), 500 μg catalase (Worthington, http://www.worthington-biochem.com), and 2 mM dithiothreitol (DTT, Solarbio), and the mixture was incubated at 30 °C for 2 h in a 15-ml gas-tight glass tube with a septum, shaking at 120 rpm, then 1 ml of gas was extracted from the headspace of the tube with a 1-ml syringe for measurement of ethylene production as previously described (Li et al., 2014). To investigate the effect of PuBZR1 on PuACO1 activity, different amounts of purified PuBZR1-His (0.2, 0.4 and 0.6 μg) were mixed with 0.2 μg of PuACO1-GST and incubated on ice for 1 h, shaking at 100 rpm. The mixture was then added to incubation buffer and incubated at 30 °C for 2 h to measure ethylene production, which was defined as the amount of ethylene produced at 30 °C in one hour.

The ACO activity of extracts from pear fruit was measured as described in Ververidis and John (1991) with a few modifications. Briefly, 1 g of fruit flesh was ground into fine powder in liquid nitrogen and suspended in 2 ml of extraction buffer containing 0.1 M Tris-HCl (pH 7.5, Sangon Biotech), 10% (v/v) glycerol, 5% polyvinylpolypyrrolidone (PVP, Solarbio), 5 mM DTT, 30 mM Na-ascorbate and 0.1 mM FeSO_4_. The suspension was centrifuged at 4 °C and 12,000 g for 10 min, and the supernatant was collected as a crude extract. To determine ACO activity, 400 μl of crude extract was incubated with 3,600 μl of a solution containing 0.1 M Tris-HCl (pH 7.5), 10% (v/v) glycerol, 1 mM ACC, 30 mM NaHCO_3_, 30 mM Na-ascorbate and 0.1 mM FeSO_4_ in a 30 °C water bath for 1 h in a 15-ml gas-tight glass tube with a septum. Ethylene production was measured to calculate the ACO activity as described above. Each experiment was repeated independently at least three times, and a Student’s *t*-test was employed to determine the statistical significance.

### Yeast One-Hybrid Assay

The *PuBZR1* and *PuERF2* CDS regions were ligated into the pGADT7 vector using the *Nde*I and *Eco*RI restriction sites. The *PuERF2* (1,979 bp upstream of the predicted translation start site), or *PuACO1* (1,941 bp upstream of the predicted translation start site) promoter fragments were cloned into the pAbAi vector using the *Kpn*I and *Xho*I restriction sites. The yeast one-hybrid (Y1H) assay was conducted using the Matchmaker™ Gold Yeast One-Hybrid Library Screening System kit (Cat. no. 630491, Clontech).

### Electrophoretic Mobility Shift Assay

For the electrophoretic mobility shift assay (EMSA), recombinant His-tagged PuBZR1 or GST-tagged PuERF2 was expressed in *E. coli* BL21 (DE3) cells and purified as described above. The biotin-labeled *PuACO1* or *PuERF2* promoter regions contained a BRRE or DRE motif as shown in Fig. 3 and Fig. 4. Corresponding unlabeled regions were used as competitors. The EMSA analysis was completed as previously described (Li et al., 2016).

### ChIP-PCR Analysis

The *PuBZR1* CDS was ligated into the pCAMBIA1307 vector as in co-IP assay. The *PuERF2* CDS was cloned into the pRI101 vector (TakaRa) to allow expression of PuERF2 as a fusion with green fluorescent protein (GFP) driven by the *CaMV 35S* promoter, using the *Kpn*I and *Eco*RI sites. The resulting *Pro35S:Myc-PuBZR1* or *Pro35S:GFP-PuERF2* constructs were transformed into pear calli, and ChIP assays were performed as previously described (Li et al., 2017) with anti-Myc or an anti-GFP antibodies. The amount of immunoprecipitated chromatin was determined by qPCR as described in Li et al. (2017). Each ChIP assay was repeated three times and the enriched DNA in each time was used as one biological replicate for qPCR. At least three biological replicates were performed and a Student’s *t*-test was employed to determine the statistical significance. Primers used are listed in Supplemental Data Set 3.

### GUS Analysis

The *PuBZR1* or *PuERF2* CDS regions were cloned into the pRI101 vector (Xiao et al., 2013) using restriction enzymes sites (*Nde*I and *Eco*RI for *PuBZR1, Nde*I and *Bam*HI for *PuERF2*) to generate the effector constructs. The reporter constructs were generated using the *PuACO1* (1,941 bp), *PuACS1a* (1,298 bp) and *PuERF2* (1,979 bp) promoter sequences cloned upstream of the GUS reporter gene in the pBI101 vector. The reporter and effector vectors were transformed into *A. tumefaciens* strain EHA105, and tobacco (*N. benthamiana*) leaves were used for co-infiltration. The co-infiltration and examination of GUS activity was performed according to Li et al. (2017). The infiltration in each assay was repeated three times as three biological replicates, and a Student’s *t*-test was employed to determine the statistical significance.

### Firefly Luciferase Complementation Imaging Assay

The *PuBZR1* or *PuACO1* CDS regions were inserted into the pCAMBIA1300-nLuc/-cLuc vectors (Chen et al., 2008) using the *Kpn*I and *Sal*I or *Kpn*I and *Pst*I restriction enzyme sites, respectively. *A. tumefaciens* strain EHA105 carrying the indicated constructs was cultured to OD_600_ 0.5 and incubated at room temperature for 3 h before being infiltrated into tobacco leaves. The EBR treatment (10 μM) was applied to *Agrobacterium*-infiltrated tobacco leaves 3 h before imaging and luciferase activity was detected as previously described (Li et al., 2017). The infiltration in each assay was repeated three times as three biological replicates.

### Agrobacterium Infiltration

To silence *PuBZR1* expression in pear fruit, the full *PuBZR1* CDS was ligated into the pRI101 vector in the reverse direction to generate the antisense *pRI101-PuBZR1-AN* construct. The recombinant construct was transformed into *A. tumefaciens* strain EHA105. The infection suspension was prepared as in Li et al. (2016). Infiltration assay was performed on pear (*P. ussuriensis* cv. Nanguo) fruit still attached to trees approximately 7 days before commercial harvest. For silencing of *PuBZR1* expression, 100 μl of the suspension was taken using a 1-ml sterile syringe and injected into fruit at a depth of 0.5 cm and 4-5 injections were performed on each fruit. Infiltrated fruit were harvested 7 days after infiltration, treated with 3 μM of EBR as above, stored at room temperature for 10 d and sampled every 5 d. One fruit was used as a biological replicate and at least three fruit were used for measurement of ethylene production or gene expression at each sampling point, and a Student’s *t*-test was employed to determine the statistical significance.

### RNA Sequencing of Apple Fruit

Control and EBR treated apple fruits sampled at day 10 (stored at room temperature for 10 d after harvest) were used for RNA sequencing (RNA-seq). RNA extracted from control or EBR treated fruits (three biological replicates for each) was used for library construction, and a total of six libraries were constructed. cDNA synthesis and library construction were performed according to previously described (Huang et al., 2014). RNA-seq was performed using an Illumina HiSeq2500 by BIOMARKER (http://www.biomarker.com.cn/). The FPKM (reads per kb per million reads) method was used to calculate the rate of differential expressed genes. The false discovery rate (FDR) was used to determine the *p*-value thresholds via multiple testing. All genes with a Log2FC (Fold Change) greater than 1.5 or *p*-value < 0.05 were selected. The Unique gene identifier (Gene ID), log2FC, FDR and annotation are indicated in Supplemental Data Set 1. The heat map for differentially expressed genes between untreated and EBR treated apple fruits was constructed using Cluster 3.0 software. All the raw data has been deposited into NCBI Sequence Read Archive (SRA) under accession number PRJNA557322.

### Accession Numbers

Sequence data from this article can be found in the Genome Database for Rosaceae (https://www.rosaceae.org) or GenBank libraries under accession numbers *PuBZR1* (MH188908), *PuERF2* (MH188911), *PuERF3* (MH188907), *PuERF106* (MH188910), *PuERF113* (MH188909), *PuACO1* (MH188913), *PuACS1a* (EF566865), *PuACS1b* (KC63252), *PuACS1-like* (XM018643584), *PuACS10* (XM009375065), *PuACS12* (XM009376269), *PuActin* (AB190176), *MdBZR1* (MDP0000306427), *MdACS1* (U89156), *MdACO1* (AF030859), *MdActin* (EB136338), *SlACO1* (EF501822), and *PaACO1* (M32692).

## SUPPLEMENTAL DATA

**Supplemental Figure S1**. Ethylene production in pear fruit treated with epibrassinolide (EBR).

**Supplemental Figure S2**. Expression of *PuBZR1* and its homologs in pear fruit treated with epibrassinolide (EBR).

**Supplemental Figure S3**. The interaction between PuBZR1 and PuACO1 proteins and their intracellular localization.

**Supplemental Figure S4**. Sequence alignment of *PuACO1* with its homologs from tomato (*Solanum lycopersicum*), apple (*Malus domestica*) and avocado (*Persea americanna*).

**Supplemental Figure S5**. ACC oxidase activity in pear fruit treated with epibrassinolide (EBR).

**Supplemental Figure S6**. Expression of *PuACSs* in pear fruit treated with epibrassinolide (EBR).

**Supplemental Figure S7**. Expression of *PuERFs* in pear fruit treated with epibrassinolide (EBR).

**Supplemental Figure S8**. PuBZR1 works upstream of PuERF2.

**Supplemental Figure S9**. PuBZR1 does not interact with PuERF2 in yeast cells.

**Supplemental Figure S10**. The interaction between PuBZR1 and PuACO1 does not affect the binding of PuBZR1 to its target promoters.

**Supplemental Figure S11**. Heat map of differentially expressed genes between untreated and EBR treated apple fruits from the RNA sequencing data.

**Supplemental Data Set 1**. Differentially expressed genes identified from RNA-seq data of apple fruit treated with or without epibrassinolide (EBR).

**Supplemental Data Set 2**. Brassinosteroids (BR)-suppressed ERF transcription factors identified from RNA-seq data of apple fruit treated with or without epibrassinolide (EBR).

**Supplemental Data Set 3**. Primers used in this study.

## ACKNOWLEDGMENTS

This work was supported by the National Key Research and Development Program of China (2018YFD1000105) and the National Natural Science Foundation of China (31722047). We thank Professor Nan Ma from China Agricultural University for kindly providing the NF-YA4-mcherry vector, and Professor Zhi Liu from Liaoning Pomology Institute for kindly providing the apple fruit samples. We also thank PlantScribe (http://www.plantscribe.com) for editing this manuscript.

## LITERATURE CITED

Alayón-Luaces P, Pagano EA, Mroginski LA, Sozzi GO (2008) Four glycoside hydrolases are differentially modulated by auxins, cytokinins, abscisic acid and gibberellic acid in apple fruit callus cultures. Plant Cell, Tissue and Organ Culture 95: 257–263

Baghel M, Nagaraja A, Srivastav M, Meena NK, Senthil Kumar M, Kumar A, Sharma RR (2019) Pleiotropic influences of brassinosteroids on fruit crops: a review. Plant Growth Regulation 87: 375–388

Barry CS, Giovannoni JJ (2007) Ethylene and Fruit Ripening. Journal of Plant Growth Regulation 26: 143–159

Chai Y-m, Zhang Q, Tian L, Li C-L, Xing Y, Qin L, Shen Y-Y (2012) Brassinosteroid is involved in strawberry fruit ripening. Plant Growth Regulation 69: 63–69

Chen H, Zou Y, Shang Y, Lin H, Wang Y, Cai R, Tang X, Zhou JM (2008) Firefly luciferase complementation imaging assay for protein-protein interactions in plants. Plant Physiol 146: 368–376

Clouse SD (2011) Brassinosteroid Signal Transduction: From Receptor Kinase Activation to Transcriptional Networks Regulating Plant Development. The Plant Cell 23: 1219–1230

Dandekar AM, Teo G, Defilippi BG, Uratsu SL, Passey AJ, Kader AA, Stow JR, Colgan RJ, James DJ (2004) Effect of down-regulation of ethylene biosynthesis on fruit flavor complex in apple fruit. Transgenic research 13: 373–384

Dong JG, Fernandez-Maculet JC, Yang SF (1992) Purification and characterization of 1-aminocyclopropane-1-carboxylate oxidase from apple fruit. Proceedings of the National Academy of Sciences 89: 9789–9793

Guo YF, Shan W, Liang SM, Wu CJ, Wei W, Chen JY, Lu WJ, Kuang JF (2019) MaBZR1/2 act as transcriptional repressors of ethylene biosynthetic genes in banana fruit. Physiol Plant 165: 555–568

Gupta A, Pal RK, Rajam MV (2013) Delayed ripening and improved fruit processing quality in tomato by RNAi-mediated silencing of three homologs of 1-aminopropane-1-carboxylate synthase gene. Journal of Plant Physiology 170: 987–995

Guy M, Kende H (1984) Conversion of 1-aminocyclopropane-1-carboxylic acid to ethylene by isolated vacuoles of Pisum sativum L. Planta 160: 281–287

Han YC, Kuang JF, Chen JY, Liu XC, Xiao YY, Fu CC, Wang JN, Wu KQ, Lu WJ (2016) Banana Transcription Factor MaERF11 Recruits Histone Deacetylase MaHDA1 and Represses the Expression of MaACO1 and Expansins during Fruit Ripening. Plant Physiol 171: 1070–1084

He J-X, Gendron JM, Sun Y, Gampala SS, Gendron N, Sun CQ, Wang Z-Y (2005) BZR1 is a transcriptional repressor with dual roles in brassinosteroid homeostasis and growth responses. Science 307: 1634–1638

He Y, Li J, Ban Q, Han S, Rao J (2018) Role of brassinosteroids in persimmon (Diospyros kaki L.) fruit ripening. Journal of Agricultural and Food Chemistry 66: 2637–2644

Huang G, Li T, Li X, Tan D, Jiang Z, Wei Y, Li J, Wang A (2014) Comparative transcriptome analysis of climacteric fruit of Chinese pear (Pyrus ussuriensis) reveals new insights into fruit ripening. PLoS One 9: e107562

Ito Y, Kitagawa M, Ihashi N, Yabe K, Kimbara J, Yasuda J, Ito H, Inakuma T, Hiroi S, Kasumi T (2008) DNA-binding specificity, transcriptional activation potential, and the rin mutation effect for the tomato fruit-ripening regulator RIN. Plant J 55: 212–223

Kende H (1993) Ethylene biosynthesis. Annual review of plant biology 44: 283–307

Kim TW, Wang ZY (2010) Brassinosteroid signal transduction from receptor kinases to transcription factors. Annu Rev Plant Biol 61: 681–704

Klee HJ, Giovannoni JJ (2011) Genetics and control of tomato fruit ripening and quality attributes. Annu Rev Genet 45: 41–59

Li J, Jin H (2007) Regulation of brassinosteroid signaling. Trends Plant Sci 12: 37–41

Li T, Jiang Z, Zhang L, Tan D, Wei Y, Yuan H, Li T, Wang A (2016) Apple (Malus domestica) MdERF2 negatively affects ethylene biosynthesis during fruit ripening by suppressing MdACS1 transcription. The Plant Journal 88: 735–748

Li T, Li X, Tan D, Jiang Z, Wei Y, Li J, Du G, Wang A (2014) Distinct expression profiles of ripening related genes in the ‘Nanguo’ pear (Pyrus ussuriensis) fruits. Scientia Horticulturae 171: 78–82

Li T, Xu Y, Zhang L, Ji Y, Tan D, Yuan H, Wang A (2017) The Jasmonate-Activated Transcription Factor MdMYC2 Regulates ETHYLENE RESPONSE FACTOR and Ethylene Biosynthetic Genes to Promote Ethylene Biosynthesis during Apple Fruit Ripening. Plant Cell 29: 1316–1334

Li XJ, Chen XJ, Guo X, Yin LL, Ahammed GJ, Xu CJ, Chen KS, Liu CC, Xia XJ, Shi K, Zhou J, Zhou YH, Yu JQ (2016) DWARF overexpression induces alteration in phytohormone homeostasis, development, architecture and carotenoid accumulation in tomato. Plant Biotechnol J 14: 1021–1033

Lv B, Tian H, Zhang F, Liu J, Lu S, Bai M, Li C, Ding Z (2018) Brassinosteroids regulate root growth by controlling reactive oxygen species homeostasis and dual effect on ethylene synthesis in Arabidopsis. PLoS Genet 14: e1007144

Osorio S, Scossa F, Fernie AR (2013) Molecular regulation of fruit ripening. Front Plant Sci 4: 198

Rocklin AM, Tierney DL, Kofman V, Brunhuber NMW, Hoffman BM, Christoffersen RE, Reich NO, Lipscomb JD, Que L (1999) Role of the nonheme Fe(II) center in the biosynthesis of the plant hormone ethylene. Proceedings of the National Academy of Sciences 96: 7905–7909

Schaffer RJ, Friel EN, Souleyre EJ, Bolitho K, Thodey K, Ledger S, Bowen JH, Ma JH, Nain B, Cohen D, Gleave AP, Crowhurst RN, Janssen BJ, Yao JL, Newcomb RD (2007) A genomics approach reveals that aroma production in apple is controlled by ethylene predominantly at the final step in each biosynthetic pathway. Plant Physiol 144: 1899–1912

Shaw J-F, Chou Y-S, Chang R-C, Yang SF (1996) Characterization of the Ferrous Ion Binding Sites of Apple 1-Aminocyclopropane-1-carboxylate Oxidase by Site-Directed Mutagenesis. Biochemical and Biophysical Research Communications 225: 697–700

Symons GM, Davies C, Shavrukov Y, Dry IB, Reid JB, Thomas MR (2006) Grapes on steroids. Brassinosteroids are involved in grape berry ripening. Plant Physiol 140: 150–158

Tatsuki M, Nakajima N, Fujii H, Shimada T, Nakano M, Hayashi K, Hayama H, Yoshioka H, Nakamura Y (2013) Increased levels of IAA are required for system 2 ethylene synthesis causing fruit softening in peach (Prunus persica L. Batsch). J Exp Bot 64: 1049–1059

Trainotti L, Tadiello A, Casadoro G (2007) The involvement of auxin in the ripening of climacteric fruits comes of age: the hormone plays a role of its own and has an intense interplay with ethylene in ripening peaches. J Exp Bot 58: 3299–3308

Vardhini BV, Rao SSR (2002) Acceleration of ripening of tomato pericarp discs by brassinosteroids. Phytochemistry 61: 843–847

Ververidis P, John P (1991) Complete recovery in vitro of ethylene-forming enzyme activity. Phytochemistry 30: 725–727

Xiao YY, Chen JY, Kuang JF, Shan W, Xie H, Jiang YM, Lu WJ (2013) Banana ethylene response factors are involved in fruit ripening through their interactions with ethylene biosynthesis genes. J Exp Bot 64: 2499–2510

Yang SF, Hoffman NE (1984) Ethylene biosynthesis and its regulation in higher plants. Annual Review of Plant Physiology 35: 155–189

Yin Y, Vafeados D, Tao Y, Yoshida S, Asami T, Chory J (2005) A new class of transcription factors mediates brassinosteroid-regulated gene expression in Arabidopsis. Cell 120: 249–259

Zaharah SS, Singh Z, Symons GM, Reid JB (2011) Role of Brassinosteroids, Ethylene, Abscisic Acid, and Indole-3-Acetic Acid in Mango Fruit Ripening. Journal of Plant Growth Regulation 31: 363–372

Zhang M, Yuan B, Leng P (2009) The role of ABA in triggering ethylene biosynthesis and ripening of tomato fruit. Journal of Experimental Botany 60: 1579–1588

Zhang S, Feng M, Chen W, Zhou X, Lu J, Wang Y, Li Y, Jiang CZ, Gan SS, Ma N, Gao J (2019) In rose, transcription factor PTM balances growth and drought survival via PIP2;1 aquaporin. Nat Plants 5: 290–299

Zhang Z, Barlow JN, Baldwin JE, Schofield CJ (1997) Metal-Catalyzed Oxidation and Mutagenesis Studies on the Iron(II) Binding Site of 1-Aminocyclopropane-1-carboxylate Oxidase. Biochemistry 36: 15999–16007

Zhu T, Tan W-R, Deng X-G, Zheng T, Zhang D-W, Lin H-H (2015) Effects of brassinosteroids on quality attributes and ethylene synthesis in postharvest tomato fruit. Postharvest Biology and Technology 100: 196–204

Zhu Z, Zhang Z, Qin G, Tian S (2010) Effects of brassinosteroids on postharvest disease and senescence of jujube fruit in storage. Postharvest Biology and Technology 56: 50–55

